# Bile Acids Regulate Accumbal Cholinergic Circuitry and Dopamine Release through TGR5 Activation

**DOI:** 10.64898/2026.01.16.699938

**Authors:** Isabella A. Roque, Shimran Sharma, Philipp Mews, Summer L. Thompson, Jordan T. Yorgason

**Author notes:** Corresponding author: Jordan Yorgason. Work performed in this location (Isabella A. Roque, Shimran Sharma, Jordan T. Yorgason).

## Abstract

**Background:** Fatty foods and alcohol (i.e., ethanol) produce strong reinforcing effects, in part by altering cholinergic interneuron (CIN) activity and tonic dopamine (DA) release within the nucleus accumbens (NAc). Ethanol and fatty foods also both stimulate hepatic and possibly local brain bile acid (BA) synthesis, which raises the possibility that BAs may act as a common upstream regulator of these substances’ shared mesolimbic effects.

**Methods:** The current study investigated whether BAs can directly alter mesolimbic activity. Electrophysiological data from acute mouse brain slices was collected to assess BA effects on NAc CIN firing, as well as on excitatory and inhibitory postsynaptic CIN inputs. Bile effects on NAc DA release and clearance rates were measured through voltammetry.

**Results:** We found that low concentrations of a 1:1 mixture of BAs cholic acid (CA) and deoxycholic acid (DCA; 1-10 μM) increased CIN firing rate, whereas high BA concentrations (1-10 mM) decreased CIN firing. We further demonstrated that BA-induced excitatory effects on CIN firing are independently mediated by at least two mechanisms: Takeda G-protein-coupled receptor 5 (TGR5) activation and suppression of inhibitory CIN currents. Additionally, our results indicate that BAs modulate inhibitory input in a complex manner, reducing frequency at low concentrations, but increasing at high concentrations, and increasing amplitude at low concentrations current amplitude, and the distribution of postsynaptic current amplitude sizes across concentrations. Finally, our voltammetry data indicate that while low BA concentrations enhance NAc DA release without affecting DA uptake, high BA concentrations robustly inhibit accumbal DA release.

**Conclusion:** Our findings provide evidence that BAs exert direct modulatory effects on neural activity in the striatum.

## 1. Introduction

Fatty foods and ethanol have strong reinforcing effects, implicating changes in mesolimbic activity. High-fat diets (HFDs) have been shown to increase cholinergic interneuron (CIN)-evoked accumbal dopamine (DA) release in rat models [1]. In parallel, ethanol has been observed to both directly increase and decrease DA release and CIN firing in the nucleus accumbens (NAc), effects that are largely dose-dependent and appear to involve multiple mechanisms [2]. Together, these findings indicate that both fatty foods and ethanol disrupt CIN-DA interactions in the NAc. Given the established connection between CIN activity, DA release in the striatum, and reward-related behavior [3], identifying a common upstream regulator for fatty food- and ethanol-induced CIN-DA effects may provide insight into the mechanisms underlying their common reinforcing properties.

Notably, both fatty foods and ethanol promote hepatic, and possibly local brain, bile acid (BA) synthesis, making BAs a potential upstream regulator of these substances’ shared striatal effects [4-7]. Consumption of 1–2 standard alcohol doses increases hepatic BA production by ∼20% and elevates systemic BA precursors 5–15-fold [5, 8]. Alcohol intake also raises cholesterol levels in the brain, providing more precursors for BA synthesis in the Central Nervous System (CNS) [4]. Similarly, HFD raises systemic BA levels by up to 50% in rats and increases levels of brain BA precursor 27-hydroxycholesterol (27-OHC) and BA-synthesizing enzyme cholesterol 27α-hydroxylase (CYP27A1) [6, 7]. While brain-derived BAs can act locally, hepatic BAs must enter systemic circulation and cross the blood-brain barrier (BBB) to influence activity in the CNS [9]. Approximately 10% of BAs exiting the intestines reach systemic circulation [10]. Bile in systemic circulation can subsequently reach the BBB and cross into the CNS either via BA transporters or passive diffusion [11, 12]. There is strong evidence indicating that hepatic BAs cross the BBB and enter the CNS. Brain and serum BA concentrations are significantly correlated, and the presence of BAs such as DCA—a secondary BA that can only be synthesized outside the CNS—has been detected in brain tissue [13, 14]. The liver produces primary BAs, like cholic acid (CA), and intestinal microbiota converts these into secondary BAs, like deoxycholic acid (DCA). In contrast, the brain only expresses enzymes for primary BA synthesis [9, 13]. Once they reach the CNS, both CA and DCA can activate BA receptors present in the brain, such as Takeda G-protein-coupled receptor 5 (TGR5), Farnesoid X receptor (FXR), and Sphingosine-1-phosphate receptor 2 (S1PR2) [9,15-16]. Importantly, there is evidence that TGR5 expression in the NAc contributes to cocaine’s rewarding effects, highlighting a putative role for BAs in regulating mesolimbic activity [16].

This study used a 1:1 mixture of CA and DCA to examine general BA effects on mesolimbic circuitry, irrespective of their physiological brain- or hepatic-derived source. We assessed BA effects on CIN firing and DA release and evaluated the role of the BA receptor TGR5 in mediating these effects. Our findings demonstrate that BAs exert significant mesolimbic effects *in vitro* and suggest that BAs contribute to fatty foods’ and ethanol’s shared appetitive effects *in vivo*.

## 2. Materials and Methods

### 2.1. Animals

Male and female mice on a C57BL/6J background (Jackson Laboratory) were kept in a 12:12-h light/dark cycle and given access to water and standard rodent chow *ad libitum*. Animal care and experimental procedures were approved by Brigham Young University Institutional Animal Care and Use Committee and were performed in accordance with the National Institutes of Health Guide for the Care and Use of Laboratory Animals.

### 2.2. Brain Slice Preparation and Electrophysiology Recordings

Fully isoflurane (Patterson Veterinary, Devens, MA) anesthetized mice (PND>30) were decapitated and brains were quickly extracted and sliced into 220 µm thick striatal coronal brain slices on a vibratome (Leica VT1000S, Vashaw Scientific, Norcross, GA) surrounded by pre-oxygenated ACSF (95% O_2_/5% CO_2_; pH≈7.4, 35°C) and 300 μM ketamine to preserve cell health. The artificial cerebrospinal fluid (ACSF) contained (in mM): KCl (2.5), NaCl (126), glucose (11), NaHCO_3_ (25), NaH_2_PO_4_ (1.2), CaCl_2_ (2.4), MgCl_2_ (1.2). After a short incubation period (∼30 minutes), slices were transferred to the recording chamber and perfused with ACSF (34°C) at a rate of ∼1.8 ml/min. Cell-attached recordings were made with glass micropipettes (2.5-6 MΩ) pulled on a P97 Pipette puller (Sutter Instruments) and subsequently filled with ACSF (cell attached recordings) or artificial intracellular solution (whole cell recordings). Intracellular solution for whole cell recordings contained (in mM): KCl (145), NaCl (8), MgCl2 (0.2), EGTA (2), HEPES (10), and Mg-ATP (2). Whole-cell recordings were performed to isolate spontaneous inhibitory postsynaptic currents (sIPSCs; in kynurenic acid) and excitatory postsynaptic currents (sEPSCs; in picrotoxin). CINs were identified based on their large size, tonic firing, and sensitivity to muscarine. Drugs were applied as specified and for at least 10 minutes to allow for stabilization. Data were filtered at the amplifier with a 4-pole high-pass Bessel filter at 2–10 kHz (Axon Multiclamp 700A; Molecular Devices, Sunnyvale CA, USA) and digitized via a NIDAQ board (PCIe-6321; National Instruments) sampled continuously at 10kHz using Axograph Software 1.7.6. (Axograph Scientific, Sydney Australia). All cell-attached experiments consisted of continuous 1 h recordings, with a 5 min baseline and sequential 10 min exposures to each CA/DCA concentration (1 µM–10 mM). Whole-cell recordings consisted of shorter 15- and 25-min recordings to prevent confounds from re-sealing of the cell membrane [17]. The following drug concentrations were bath applied for electrophysiology and slice voltammetry experiments where specified: CA/DCA (1 µM - 10 mM; Millipore Sigma), 5-chloro-2-(ethylsulfonyl)-4-pyrimidinecarboxylic acid 3-methylphenyl ester (SBI-115; 10 µM; Cayman Chemical), kynurenic acid (KA, 2 mM; Millipore Sigma), picrotoxin (100 µM; Millipore Sigma).

### 2.3. Carbon Fiber Electrode Preparation and Voltammetry Recordings

Slices were prepared as described above, transferred to the recording chamber, and perfused with aCSF (34°C) at a rate of ∼1.8 ml/min. Fast scan cyclic voltammetry recordings were performed and analyzed using Demon Voltammetry and Analysis software [18]. The carbon fiber electrodes used for voltammetry experiments were manufactured in-house. Carbon fiber electrodes (7 μm, Thornel T-650, Cytec, Woodland Park, NJ) were aspirated into borosilicate glass capillaries (TW150, World Precision Instruments, Sarasota, FL) and pulled on a pipette puller and carbon fibers were cut to a length of 100-150 μm from the tip of the glass tube. Stimulating electrodes were prepared by pulling a glass micropipette and filled with ACSF. DA release was evoked in the NAc with a 1 pulse (30-100 μA; 0.5 ms) electrical stimulus (1/min) and detected via fast-scan cyclic voltammetry (100 ms cycle, 400 V/s). Concentration response curve experiments had an initial baseline (BL) stabilization period (∼18-20 min), followed by 10 min exposures to CA/DCA in increasing concentrations (1 µM–10 mM). For single concentration experiments, a similar BL stabilization period was used, followed by a ∼20 min exposure to 1-10 µM CA/DCA.

### 2.4. Statistical Analysis

Raw patch-clamp (Axograph) and voltammetry (Demon) data were exported to Excel for CIN firing/current rate, CIN firing variance and DA release averaging. Averages were normalized against respective BLs and transferred to GraphPad Prism 5 (GraphPad, San Diego, CA) or NCSS 8 (NCSS LLC, Kaysville UT) for statistical analysis. Since amplitude is an event-level property, sIPSCS/sEPSC current amplitude values were transferred directly from Excel to GraphPad without averaging. Voltammetry mediation and OLS regression were run in Python.

Mediation analysis was conducted to evaluate whether BA effects on DA release were (1) direct, independent of CIN activity, or (2) indirect, via modulation of CIN firing rates. Regression coefficients in this analysis represented average bile effects on CIN firing rates and DA release per unit increase in bile concentration. The coefficients were defined as follows:

▪ a: bile effects on CIN firing rates
▪ b: CIN firing rate effects on DA release, controlling for bile
▪ c′: direct bile effects on DA release, controlling for CIN firing
▪ a × b: indirect, CIN-mediated bile effects on DA release
▪ c = c′ + a × b: total bile effects on DA release

The predicted lines in Fig. 5d represent model-based predicted values derived from the mediation analysis. All remaining statistical analyses were performed in GraphPad. One- and two-way ANOVA and two-tailed unpaired t-tests were used, with Dunnett’s test for planned comparisons, or Tukey’s HSD or Bonferroni post-hoc analysis where stated. Outliers were identified using GraphPad’s outlier calculator. Significance levels are indicated on graphs with asterisks *,**,*** and correspond to significance levels *p*<0.05, 0.01 and 0.001, respectively. Figures were constructed with Prism 5 (GraphPad, San Diego, CA), Python and Adobe Illustrator software.

## 3. Results

### 3.1. Concentration-Dependent Bile Effects on CIN Firing Rate and Variance

Bath application of varying BA concentrations had a dose-dependent effect on CIN firing rate and variance. Traces displaying CIN firing patterns under each 1:1 CA/DCA mixture concentration are shown (**Fig. 1a**). Cholinergic interneuron firing rate increased by 178 ±24% with exposure to 1 μM CA/DCA and by 171±25% with exposure to 10 μM CA/DCA compared to BL (**Fig. 1b**; one-way repeated-measures ANOVA, F_6,47_=16.93, p<0.0001; Dunnett posthoc p<0.05 for BL vs. 1 μM and 10 μM CA/DCA). Conversely, CIN firing rate decreased by 90±8% with exposure to 10 mM CA/DCA (**Fig. 1b**; Dunnett post hoc p<0.05 for BL vs. 10 mM CA/DCA). Finally, 100 μM and 1 mM CA/DCA did not significantly influence CIN firing rate compared to BL (**Fig 1b**; Dunnett post hoc p>0.05 for BL vs. 100 μM and 1 mM CA/DCA). Cholinergic interneuron firing variance increased by 275±75% with exposure to 10 mM CA/DCA, but was not significantly influenced by other CA/DCA concentrations (**Fig. 1c**; one-way ANOVA, F_6,47_=4.869, p=0.0013; Dunnett post hoc p<0.05 for BL vs. 10 mM bile and p>0.05 for BL vs. 1, 10, 100 μM and 1 mM bile).

**Fig. 1.**
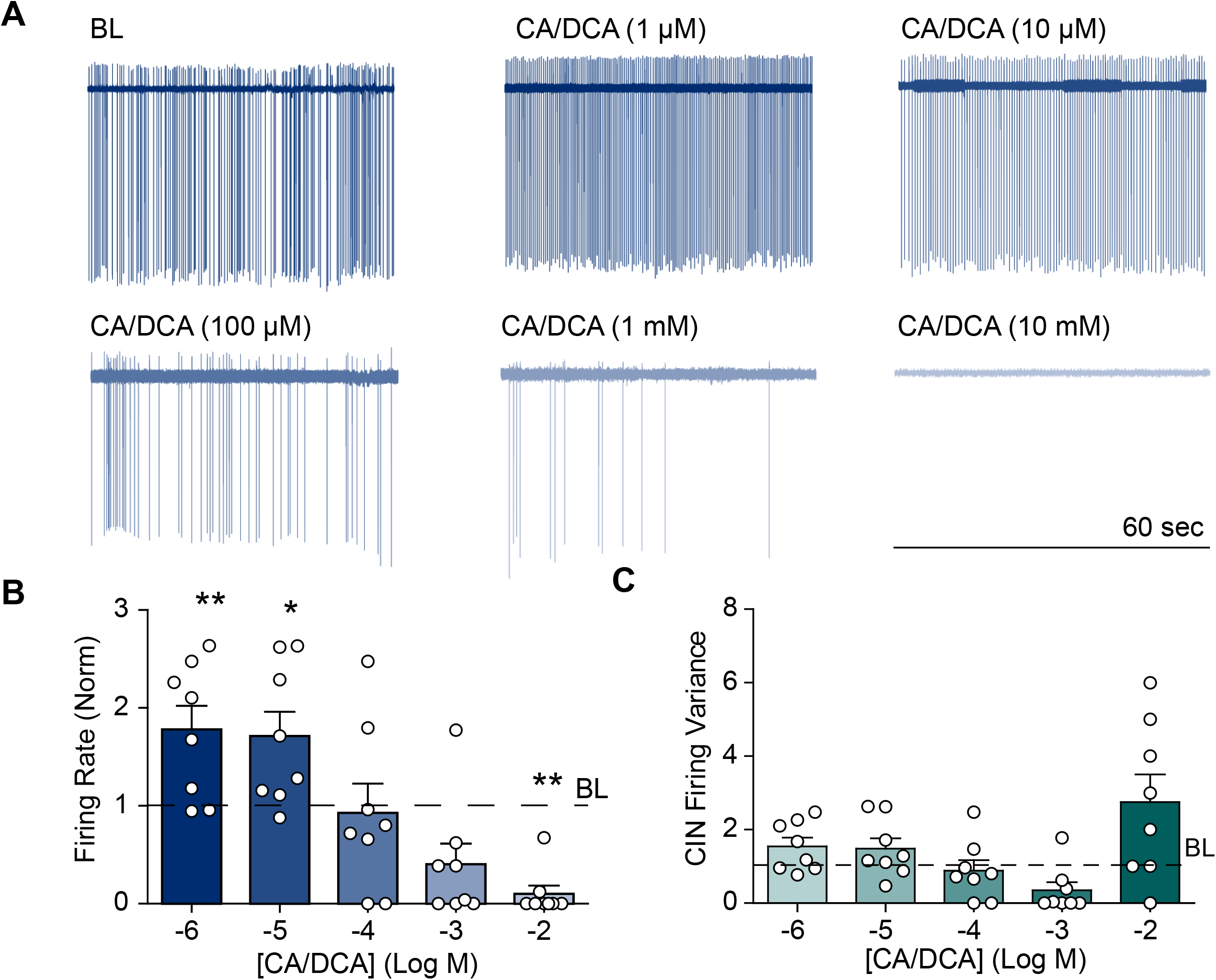
BAs exhibit concentration-dependent effects on CIN firing rate and variance. **(a)** Sample CIN firing traces during BL and 1 μM-10 mM CA/DCA conditions. **(b)** 1 μM and 10 μM CA/DCA significantly increased, 10 mM CA/DCA significantly decreased, and 100 μM and 1 mM CA/DCA did not significantly alter CIN firing rate compared to BL. **(c)** Only 10 mM CA/DCA significantly altered CIN firing variance, increasing variance significantly above BL. Statistical significance represented as * (p < 0.05), ** (p < 0.01).

### 3.2. Bile Effects on CIN Firing Rate with Exposure to TGR5 Antagonist SBI-115

Treating brain slices with a TGR5 BA receptor antagonist (SBI-115; 10 µM) altered concentration-dependent bile effects on CIN firing rate and variance (**Fig. 2**). Cholinergic firing traces under each experimental CA/DCA concentration in SBI-115-treated brain slices are shown (**Fig. 2a**). Low CA/DCA concentrations (1 μM and 10 μM) did not have a significant effect on CIN firing rate in SBI-115-treated brain slices (**Fig. 2b**; one-way repeated-measures ANOVA, F_6,42_=5.418, p=0.0008; Dunnett post hoc p>0.05 for BL vs. 1 and 10 μM CA/DCA). Intermediate and high CA/DCA concentrations (100 μM, 1 mM and 10 mM) all significantly decreased CIN firing rate by 76±13%, 77±13% and 74±25%, respectively, in SBI-115-treated brain slices compared to BL (**Fig. 2b**; Dunnett post hoc p<0.05 for BL vs. 100 μM, 1 mM and 10 mM CA/DCA). None of the experimental CA/DCA concentrations significantly altered CIN firing variance in SBI-115-treated brain slices compared to BL (**Fig. 2c**; one-way ANOVA, F_6,47_=0.3353, p=0.8887; Dunnett post hoc p>0.05 for BL vs. 1, 10, 100 μM and 1, 10 mM CA/DCA).

**Fig. 2.**
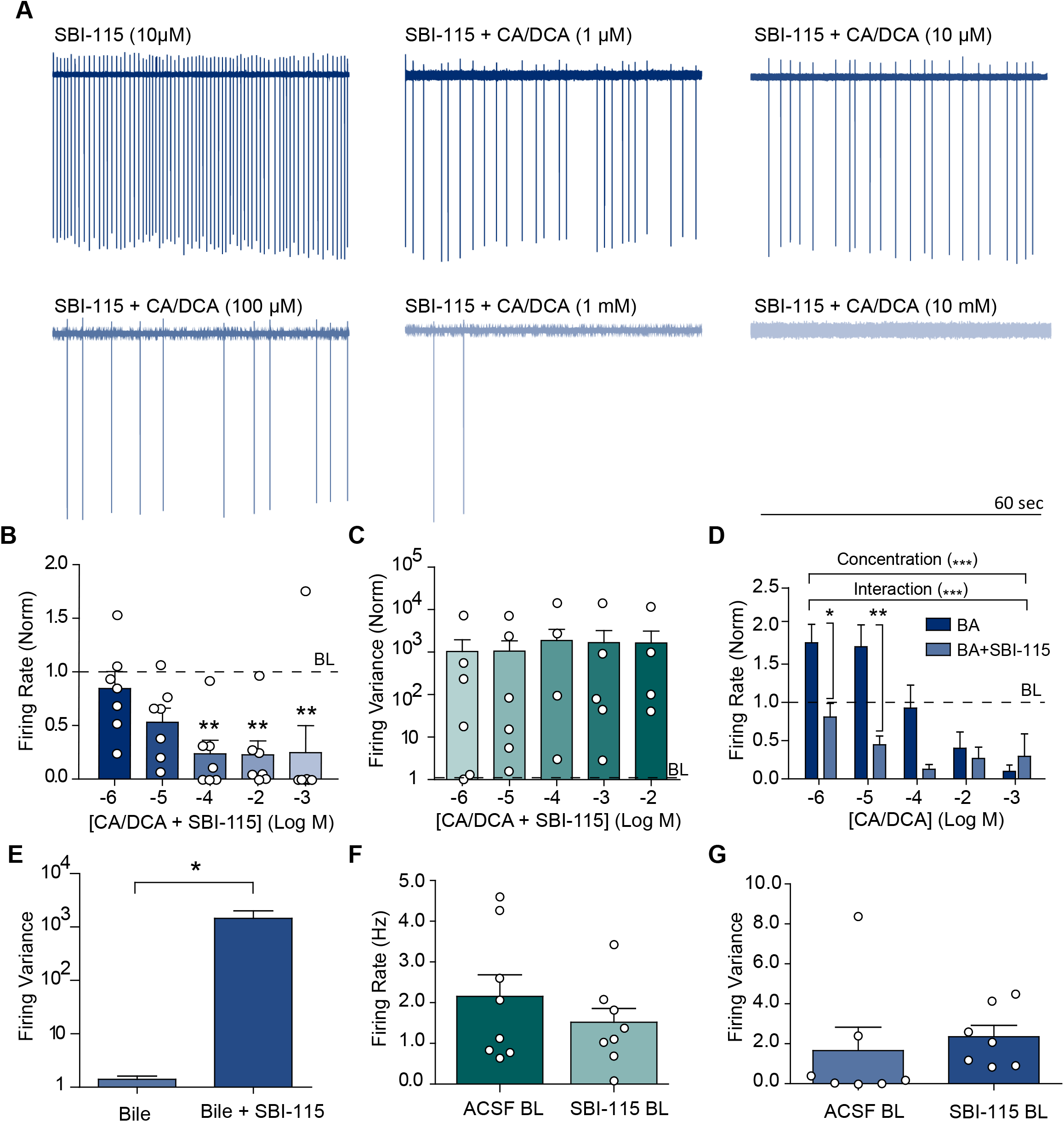
Treating brain slices with TGR5 antagonist SBI-115 alters bile effects on CIN firing rate and variance. **(a)** Sample CIN firing traces during BL and 1 μM-10 mM CA/DCA conditions in SBI-115-treated brain slices. **(b)** In SBI-115-treated brain slices, 100μM, 1mM and 10mM CA/DCA significantly decreased, and 1 and 10μM CA/DCA did not significantly alter CIN firing rate compared to BL. **(c)** None of the experimental CA/DCA concentrations significantly altered CIN firing variance in SBI-115-treated brain slices. **(d)** Bath application of SBI-115 significantly altered 1μM and 10μM CA/DCA effects on CIN firing rate, but not 100μM, 1mM and 10mM CA/DCA effects on CIN firing rate. **(e)** Addition of SBI-115 to varying CA/DCA concentration conditions significantly increased overall CIN firing variance. **(f)** CIN firing rate in standard ACSF BL conditions was not significantly different from CIN firing rate in SBI-115-treated BL ACSF conditions. **(g)** CIN firing variance was also not significantly different between standard ACSF BL conditions and SBI-115-treated BL conditions. Statistical significance represented as * (p < 0.05), ** (p < 0.01), *** (p < 0.001).

Comparison of BA effects in the presence and absence of SBI-115 showed that bath application of SBI-115 decreased CIN firing rate responses to 1 μM and 10 μM CA/DCA concentrations by 60±12%, but did not significantly alter CIN firing rate responses to 100 μM, 1 mM and 10 mM CA/DCA concentrations (**Fig. 2d**; two-way repeated-measures ANOVA (bile vs. bile + SBI-115); concentration: F_6,48_=14.59, p<0.0001; interaction: F_6,48_=5.887, p=0.0006; Dunnett’s post hoc, p<0.05 for 1 μM CA/DCA, p<0.001 for 10 μM CA/DCA and p>0.05 for 100 μM, 1 mM and 10 mM CA/DCA). Although SBI-115 did not alter CIN firing variance at individual CA/DCA concentrations, when its effects across all BA conditions were pooled, SBI-115 significantly increased overall variance compared to variance during BA conditions sans SBI-115 (**Fig. 2e**; two-tailed unpaired t-test, t_78_=2.631, p=0.0103). To test for potential independent SBI-115 effects, CIN firing rate and variance in standard ACSF BL conditions was compared to CIN firing rate and variance in SBI-115-treated ACSF BL conditions and no significant differences were found for either firing rate or variance (**Fig. 2f;** firing rate, two-tailed unpaired t-test, t_14_=1.009, p=0.3301; **Fig. 2g;** firing variance, two-tailed unpaired t-test, t_12_=0.5325, p=0.6041).

### 3.3. Bile Effects on CIN Inhibitory Current Rate and Amplitude

To further investigate how BAs influence CIN firing properties, we conducted whole-cell experiments assessing BA impact on CIN sIPSC rate and amplitude (**Fig. 3**). BAs had a concentration-dependent effect on CIN sIPSC rate and amplitude. Representative traces displaying CIN sIPSCs across varying CA/DCA concentrations are shown for BL, 10 μM, 100 μM, and 1 mM CA/DCA conditions (**Fig. 3a**). sIPSC rate was not significantly affected by 100 μM CA/DCA, a concentration that also had not previously affected firing rate or variance (**Fig. 1b-c**), yet was significantly decreased with exposure to 10 μM CA/DCA compared to the 1 mM KA BL (**Fig. 3b**; one-way ANOVA, F_2,17_= 5.083, p= 0.0186, Dunnett post hoc p<0.05 for BL vs. 10 μM CA/DCA and p>0.05 for BL vs. 100 μM CA/DCA), a dose that had increased firing rate (**Fig. 1b**). sIPSC rate was also not significantly affected by 1 mM CA/DCA (**Fig. 3c**; two-tailed unpaired t-test, t_7_=1.612, p=0.151), a dose that also had not affected firing rate or variance (**Fig. 1b-c**).

**Fig. 3.**
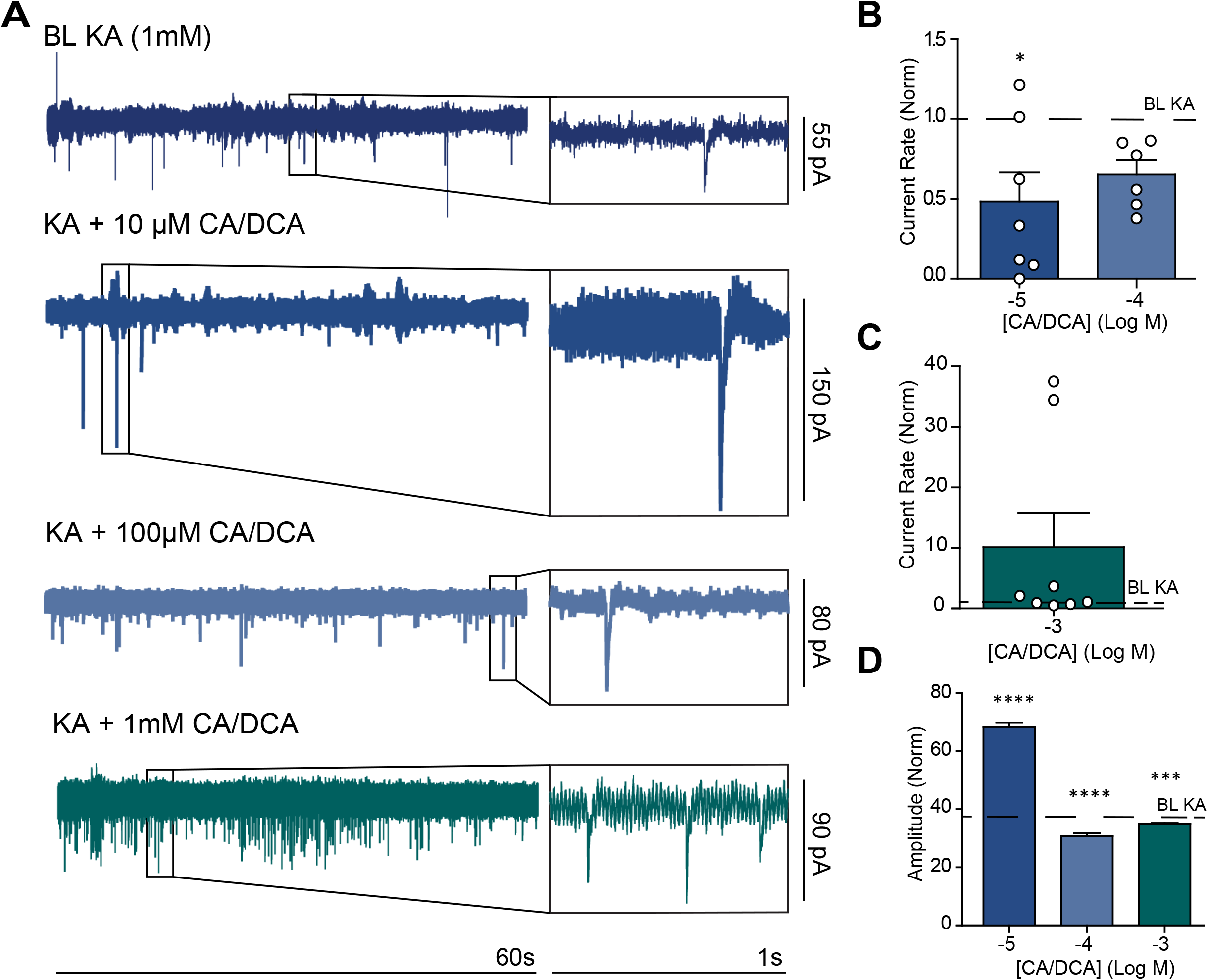
BAs have a concentration-dependent effect on CIN inhibitory current rate and amplitude. **(a)** Sample CIN firing traces with 1s expanded views of whole-cell currents during a 1mM KA BL condition, and with the addition of 10 μM-1 mM CA/DCA to BL conditions. **(b)** 10 μM CA/DCA significantly decreased and 100 μM CA/DCA did not significantly influence CIN inhibitory current rate. **(c)** 1 mM CA/DCA did not have a significant or consistent effect on CIN inhibitory current rate. **(d)** 10 μM CA/DCA significantly increases, and 100 μM and 1mM CA/DCA significantly decrease average CIN inhibitory current amplitude. Statistical significance represented as * (p < 0.05), ** (p < 0.01), *** (p < 0.001), **** (p < 0.0001).

Additionally, BAs exerted a dose-dependent effect on sIPSC amplitude. Exposure to 10 μM CA/DCA increased, whereas 100 μM and 1 mM CA/DCA decreased sIPSC amplitude (**Fig. 3d**; one-way ANOVA, F_3, 8233_= 281.7, p<0.0001, Dunnett post hoc p<0.0001 for BL vs. 10 μM CA/DCA, p<0.0001 for BL vs. 100 μM CA/DCA, and p<0.0002 for BL vs. 1 mM CA/DCA). As expected, 10 μM CA/DCA increased the frequency of high-amplitude inhibitory currents (one-way ANOVA, F_4,1521_= 3.88, p=0.0039, Dunnett post hoc p<0.05 for BL vs. 10 μM CA/DCA). Interestingly, despite decreasing the average sIPSC amplitude, 1 mM CA/DCA increased the frequency of high-amplitude sIPSCs (Dunnett post hoc p<0.05 for BL vs. 1 mM CA/DCA). In contrast, the 100 μM CA/DCA concentration did not significantly alter the frequency distribution of sIPSC amplitudes (Dunnett post hoc p>0.05 for BL vs. 100 μM CA/DCA).

### 3.4. Bile Effects on Inhibitory and Excitatory Inputs on CINs and role of TGR5

To extend our analysis of TGR5 involvement in mediating bile effects on CIN firing, we examined BA effects on sIPSCs in the presence of TGR5 antagonist SBI-115 (**Fig. 4a,b**). 10 μM CA/DCA continued to suppress sIPSCs in the presence of SBI-115 (**Fig. 4b**; two-tailed unpaired t-test, t_5_=2.224, p=0.0384), suggesting that TGR5 is not involved in BA regulation of inhibitory CIN input frequency. We next examined 10 μM BA effects on CIN excitatory inputs to determine whether a BA-induced reduction in sEPSCs could account for the effects of low BA concentrations on CIN firing (**Fig. 4c,d**). 10 μM CA/DCA did not significantly affect CIN sEPSC frequency; therefore, BA effects on excitatory CIN inputs were not explored further (**Fig. 4d**; two-tailed unpaired t-test, t_4_=1.031, p=0.1804).

**Fig. 4.**
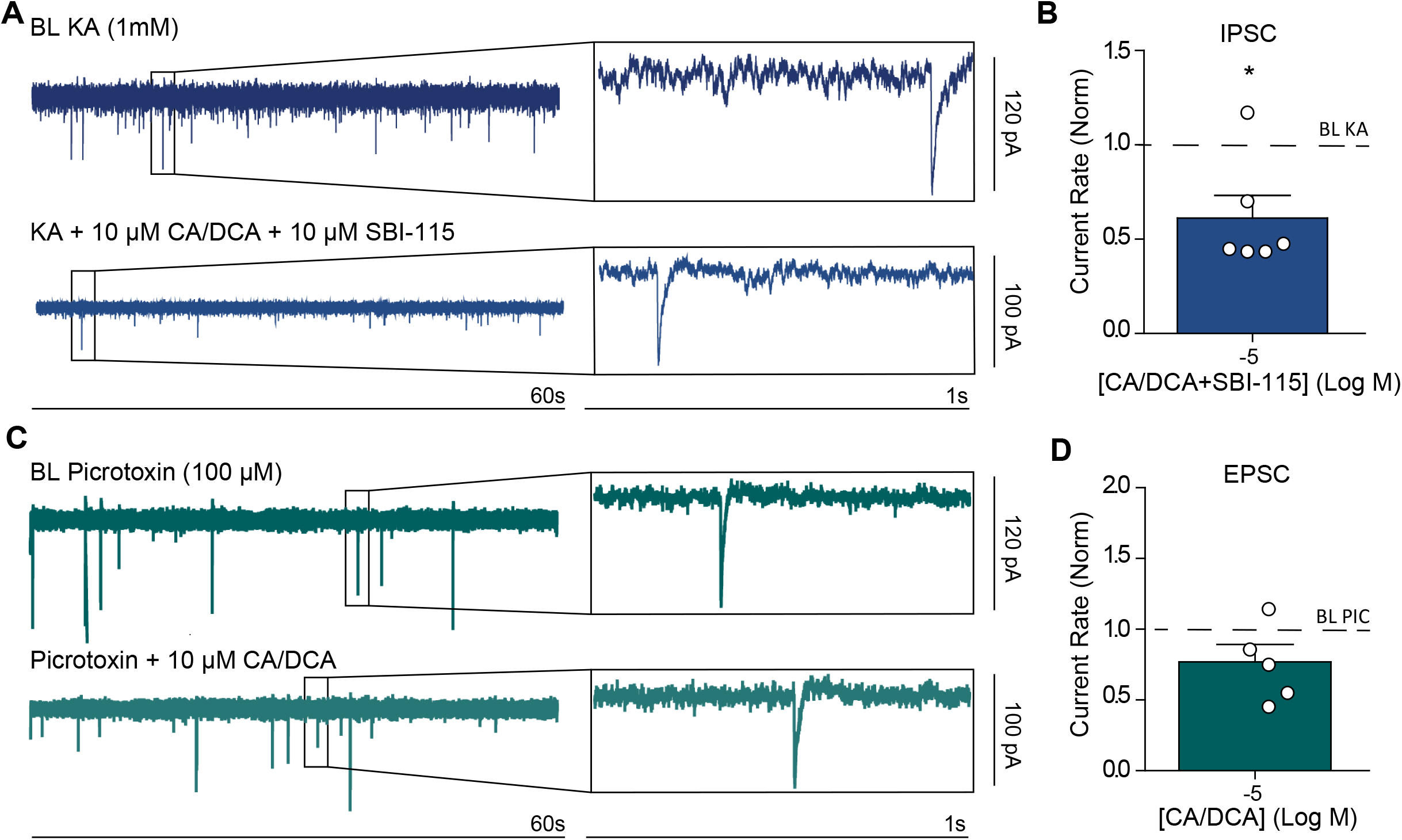
Bath application of 10 μM CA/DCA continues to suppress CIN inhibitory current rate during exposure to SBI-115 and does not significantly influence excitatory CIN current rate. **(a)** Sample CIN firing traces with enlarged 1s views of whole-cell currents at a 1mM KA BL condition and during exposure to 10 μM CA/DCA and 10 μM SBI-115 in addition to BL conditions. **(b)** During exposure to SBI-115, 10 μM CA/DCA continues to suppress CIN inhibitory current rate. **(c)** Representative CIN firing traces with 1s expanded views of whole-cell currents recorded at a 100 μM picrotoxin BL condition and during exposure to 10 μM CA/DCA in addition to BL conditions. 10 μM CA/DCA does not have a significant effect on CIN excitatory current rate. Statistical significance represented as * (p < 0.05).

### 3.5. Direct and CIN-Mediated Bile Effects on DA Release in the NAc

Given the established role of cholinergic inputs in regulating DA terminals [2, 18-20], voltammetry was used to examine whether BAs might exert indirect effects on DA transmission via modulation of CIN firing rates (**Fig. 5**). Current traces showing DA release in the presence of varying CA/DCA concentrations (1 μM-10 mM) are shown (**Fig. 5a-b**). Average DA release was not significantly altered by 1 μM-1 mM CA/DCA concentrations during continuous recordings with sequential 10-min exposures to increasing bile concentrations (**Fig. 5c**; one-way repeated-measures ANOVA, F_6,35_=10.75, p<0.0001; Dunnett post hoc p>0.05 for BL vs. 1, 10, 100 μM and 1 mM CA/DCA). However, average DA release during these continuous recordings was decreased by 96±3% with exposure to 10 mM CA/DCA (Fig. 5c; Dunnett post hoc p<0.05 for BL vs. 10 mM), with no restoration of DA release during the wash condition (Dunnett post hoc p<0.05 for BL vs. wash).

**Fig. 5.**
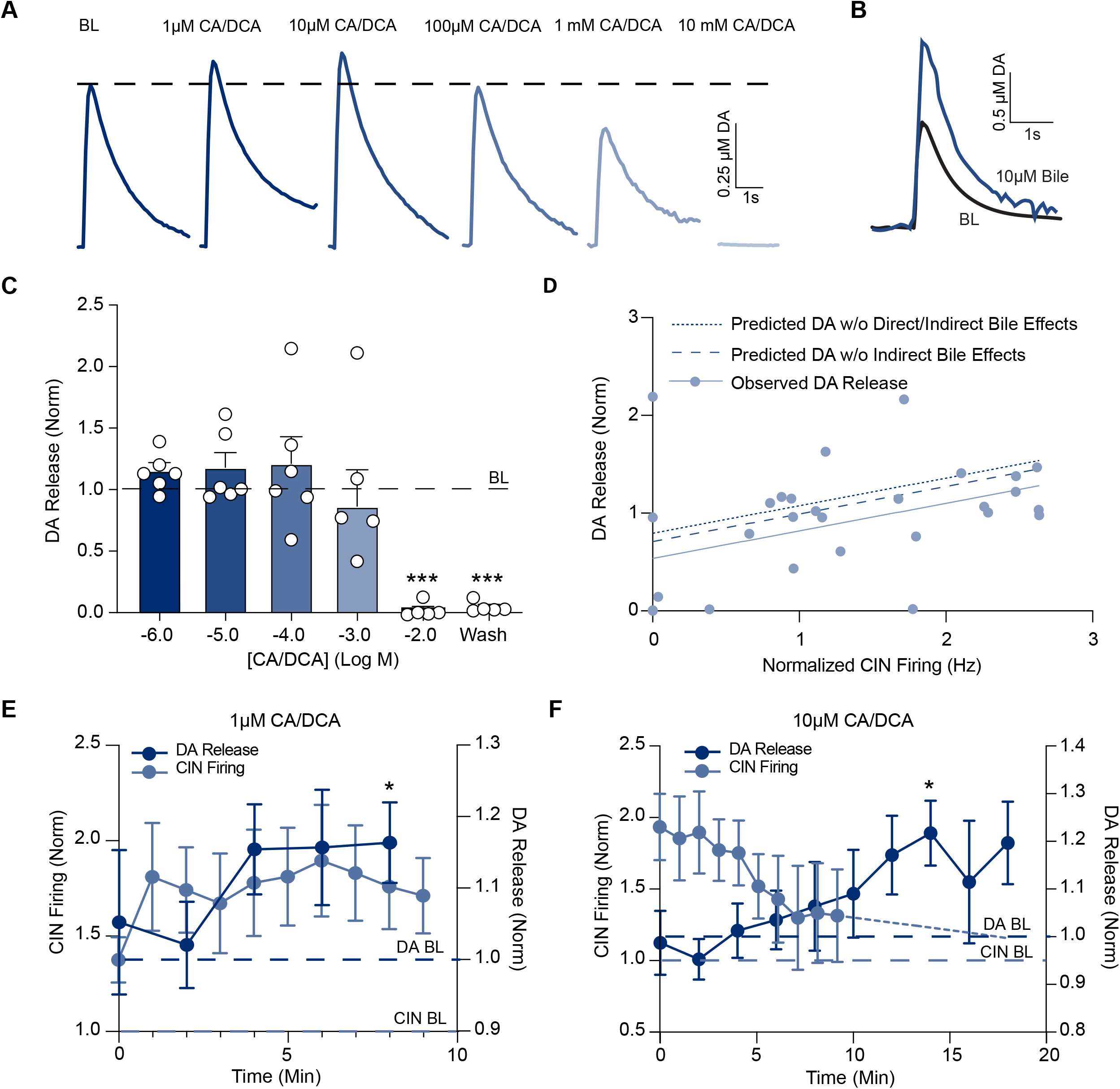
Bile Exerts Direct and CIN-Mediated Effects on Accumbal DA Release. **(a)** Sample current traces of DA release during BL, and 1 μM-10 mM CA/DCA conditions. **(b)** Representative current trace at BL and after 14 min of exposure to 10 μM CA/DCA. **(c)** 10 mM CA/DCA significantly decreased DA release, but no other experimental CA/DCA concentration conditions significantly altered average DA release. DA release was not restored during the wash condition. **(d)** Bile’s effects on average DA release at 10 mM CA/DCA are largely indirect (CIN-mediated). Dashed lines indicate predicted DA release after removing indirect (I) and both direct and indirect (D/I) bile effects. **(e)** DA release exhibits a transient increase at 8 min following bath application of 1 μM CA/DCA. **(f)** Following 10 μM CA/DCA application, DA release transiently rises at 14 min. Dashed line extending from the solid CIN firing trace represents the linear regression projection of firing beyond 10 min. Statistical significance represented as * (p < 0.05), ** (p < 0.01), *** (p < 0.001).

Mediation analysis was conducted to determine whether BA effects on average DA release at 10 mM CA/DCA were primarily direct or CIN-mediated, with the Sobel test used to assess the significance of the indirect CIN-mediated effect. We found that the inhibition of DA release that occurs with exposure to 10 mM CA/DCA is largely mediated by suppression of CIN activity, accounting for 66.9% of bile’s effect on DA release at this concentration (indirect effect: a*b = –0.1726, Sobel Z = –2.749, p = 0.006; CIN suppression: a = –0.4568, p < 0.001). Bile’s direct effects on average DA release at 10 mM CA/DCA was nonsignificant (c′ = –0.0852, p = 0.272). In **Fig. 5d**, the solid line represents observed average DA release across varying CIN firing levels, while the dashed lines show predicted DA release after accounting for the indirect (CIN-mediated) and total bile effects (total effect: c = – 0.2578, p < 0.001).

In contrast to the continuous voltammetry recordings, shorter recordings involving exposure to a single CA/DCA concentration reduced variability introduced during longer recordings and showed consistent, transient ∼20% increases in DA release with exposure to 1–10 μM CA/DCA (one-way ANOVA, F_13,52_ = 3.096, p = 0.0019). Exposure to 1 μM CA/DCA resulted in transient DA release elevations at 8 min (**Fig. 5e**; one-sample two-tailed t-test, t_5_ = 2.90, p = 0.0337), and exposure to 10 μM CA/DCA enhanced DA release at 14 min after drug application compared to BL (**Fig. 5b, f**; one-sample two-tailed t-test, t4 = 2.90, p = 0.0335). Overall, these findings are consistent with BA effects on CIN firing, as DA release showed slight increases at low BA concentrations (1–10 μM) and robust decreases at higher concentrations (>1 mM).

Mediation analysis was also conducted to determine whether significant BA effects on DA release at 1–10 μM CA/DCA were direct or CIN-mediated. Time-lagged Pearson correlations were used to account for the expected causal delay of CIN firing on DA release. During exposure to 1 μM CA/DCA, CIN firing at preceding time points strongly predicted DA release at 8 min, producing a significant mediated effect (ab = 0.036, Sobel Z = 2.79, p = 0.0053), whereas direct BA effects on DA release (c′ = −0.0140, p = 0.3020) and the total effect (c = 0.0216, p = 0.1913) at this concentration were nonsignificant. In contrast, exposure to 10 μM CA/DCA resulted in a nonsignificant CIN-mediated contribution (ab = –0.009, Sobel Z = –1.41, p = 0.158), an almost significant direct effect (c′ = 0.0249, p = 0.056), and a significant total BA effect on DA release (c = 0.0163, p = 0.0032).

DA release showed no significant transient alterations with exposure to 100 μM CA/DCA (one-sample two-tailed t-tests, t_5_ = 0.88–1.33, p = 0.242–0.421). Likewise, time-lagged mediation analysis revealed no evidence for CIN-mediated or direct bile effects on DA at this bile concentration (indirect effect a*b = –0.019, Sobel Z=-1.009, p = 0.313; direct effect c′ = 0.021, p = 0.463; total effect c = 0.003, p = 0.716). DA release also did not exhibit any transient alterations with exposure to 1 mM CA/DCA (one-sample two-tailed t-tests, t_5_ = –0.56–1.00, p = 0.365– 0.914).

### 3.6. Bile Effects on NAc DA Release and Clearance Rate

Obeticholic acid (OCA, synthetic BA) and lithocholic acid (LCA, secondary BA) have been reported to bind dopamine transporters (DATs) non-competitively [21, 22]. Given the established link between BAs and extracellular DA dynamics, we examined potential CA/DCA effects on DA clearance (tau), as well as downward (DV) and upward (UV) velocities of DA signals, at varying CA/DCA concentrations (1–100 μM; **Fig. 6**). Raw example traces (normalized to peak height) and subsequent statistical analyses revealed no significant changes in tau (**Fig. 6a,b;** one-way ANOVA, F_3,15_ = 0.3122, p = 0.8163) or DV (**Fig. 6c**; one-way ANOVA, F_3,15_ = 1.246, p = 0.3281). UV alterations, which is more of a measure of release than uptake, were also nonsignificant, but showed a trend toward significant increases with exposure to 1–10 μM CA/DCA (**Fig. 6d**; one-way ANOVA, F_3,15_ = 3.094, p = 0.0588), consistent with the enhanced DA release observed at these concentrations from **Fig. 5**. These results indicate that CA/DCA selectively modulates specific aspects of DA dynamics, such as release and UV, without affecting other properties, including tau and downward velocity (DV).

**Fig. 6.**
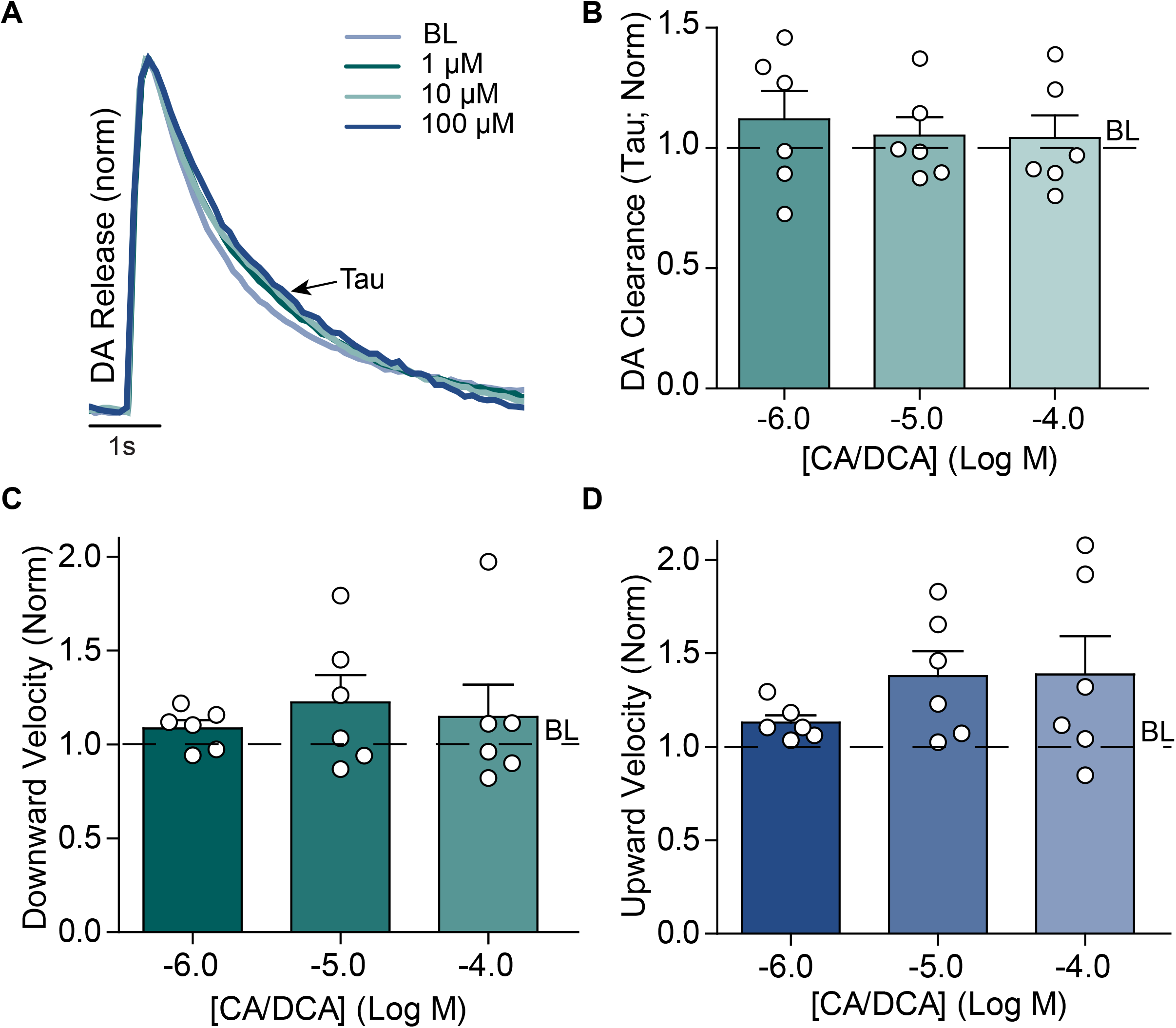
Bile (1-100 μM CA/DCA) does not affect DA clearance (tau) rates. **(a)** Overlay of sample voltammetry-based DA traces at varying CA/DCA concentrations, aligned at peak amplitude. **(b)** No significant changes were observed in DA clearance (tau) from the synaptic cleft across 1-100 μM CA/DCA concentrations. **(c)** DV, representing DAT-mediated DA clearance from the synaptic cleft, was also not significantly altered by 1-100 µM CA/DCA. **(d)** UV, reflecting DA release and helping distinguish release from clearance, showed no significant changes.

## 4. Discussion

The present study demonstrates that BAs exert potent, previously unexplored effects on mesolimbic circuitry. While low BA concentrations robustly increase CIN firing rates and weakly increase DA release, high concentrations inhibit both CIN firing and DA release, likely via membrane-destabilizing effects. TGR5 contributes to some of the excitatory CIN effects observed at low CA/DCA concentrations but does not appear to mediate the inhibitory effects seen at high CA/DCA concentrations. Notably, reductions in sIPSC frequency were not prevented by TGR5 antagonism and sEPSC frequency was unaffected with exposure to 10 μM CA/DCA, suggesting that additional receptor mechanisms may contribute to CIN excitation at this concentration. DA terminal activity closely mirrored CIN responses, indicating that bile-mediated enhancement of DA release is at least partially attributable to effects on CIN firing and that modulation of CINs is sufficient to influence downstream circuits. Finally, no direct BA effects on tau were observed, suggesting that only specific BAs (i.e., OCA and LCA) may influence DA clearance rates.

### 4.1. Concentration-Dependent Bile Effects on CIN Firing Rate and Variance

Estimates of total brain BA concentrations depend on both peripheral BA influx and potential local BA synthesis within the brain. Under normal physiological conditions, BAs produced in the liver contribute approximately 0.2–0.4 µM to total brain BA levels, while brain-resident BA-synthesizing enzymes may contribute an additional 0.05–0.1 µM, resulting in a net estimated brain BA concentration of ∼0.25–0.5 µM [17, 23]. Following moderate ethanol intake (1–2 standard drinks), peripheral BA contributions may rise to ∼0.5–2 µM and local BA synthesis— potentially via increased expression of enzymes like CYP46A1—may increase to ∼0.2–0.5 µM, yielding an estimated total brain concentration of ∼0.7–2.5 µM [5, 12]. Similarly, HFD-induced increases in hepatic BA secretion are estimated to elevate peripheral BA contributions to brain concentrations to approximately 0.3–0.75 µM, with local synthesis potentially adding another 0.1–0.3 µM. This results in total brain BA levels in the range of ∼0.4–1 µM [6, 7].

The current study uses a 1:1 CA/DCA mixture to assess general BA effects on striatal activity, regardless of their possible brain-derived or hepatic origin. Our results show that low concentrations (1–10 μM) of the 1:1 CA/DCA mixture increase CIN firing rate, whereas high concentrations (10 mM) suppress CIN activity. Our BL brain BA estimates (∼0.2–0.4 µM) and post-ethanol/HFD brain BA estimates (∼0.4–2.5 µM) indicate that ethanol and fatty food consumption can elevate brain BA levels into a concentration range (∼1 µM) that, in our *in vitro* experiments, is sufficient to increase CIN activity. Heightened CIN activity in the NAc could, in turn, promote DA release, providing a potential mechanism by which BAs can influence mesolimbic signaling and reinforce hedonic feeding and ethanol consumption [2]. Our results also demonstrate that exposure to 10 mM CA/DCA significantly decreases CIN firing rate. This inhibitory effect at moderate to high BA concentrations may reflect genuine suppression of CIN activity or could result from a compromised cell membrane potentially leading to cell death, especially at the highest concentration [24]. Additionally, we show that BAs exert a concentration-dependent effect on CIN firing variance. Specifically, exposure to 10 mM CA/DCA significantly increases CIN firing variance, which may reflect irregular firing patterns—such as sparse and burst firing—that sometimes precede neuronal death in patch clamp experiments.

### 4.2. Bile Effects on CIN Firing Rate with Exposure to TGR5 Antagonist SBI-115

The TGR5 receptor is activated by several BAs, including CA and DCA [25], and is present in neurons and astrocytes within the CNS [15]. Furthermore, histological data available on the Allen Mouse Brain Atlas confirm TGR5 expression in the NAc of adult mouse brain slices [26], and behavioral studies in TGR5 knockout mouse models demonstrate that this receptor is necessary for cocaine reward [16]. In the present study, we investigated whether TGR5 activation contributes to BA effects on accumbal CIN firing rate. Bath application of TGR5 antagonist SBI-115 abolished the excitatory effects of low (1–10 μM) CA/DCA concentrations on CIN firing but had no effect on the inhibitory response observed at high (10 mM) CA/DCA concentrations.

Our findings demonstrate that low, physiologically relevant BA concentrations enhance CIN activity through TGR5 activation, whereas the inhibitory effects observed at high, supraphysiological concentrations are mediated by TGR5-independent mechanisms. Supporting this, SBI-115 alone did not alter baseline CIN firing, indicating that it has no independent effect but instead selectively blocks TGR5-mediated excitation at low BA concentrations. In contrast, the TGR5-independent nature of high BA concentration effects supports the interpretation that supraphysiological BA levels act through nonspecific mechanisms, such as compromising membrane integrity, which leads to neuronal dysfunction or cell death and manifests as apparent inhibition of CIN activity. The lack of firing rate recovery during washout—regardless of SBI-115 presence—further reinforces the possibility of membrane disruption and irreversible cell damage at high CA/DCA concentrations (>1 mM).

Our results additionally show that although bath application of SBI-115 did not affect BL CIN firing variance, SBI-115 exposure selectively increased variance in response to all experimental BA concentrations (see Fig. 2c,e), suggesting that TGR5 signaling acts as a key stabilizer of CIN firing during elevated BA exposure. Cholinergic interneurons normally exhibit irregular firing patterns, and previous studies have shown that activation of specific channels, such as the small-conductance calcium-activated potassium channel (SK) and the hyperpolarization-activated cyclic nucleotide-gated channel (HCN, Ih), can reduce variability in CIN activity [27, 28]. Activation of the TGR5 receptor may trigger many of the same downstream mechanisms that stabilize CIN firing following SK and HCN channel activation. This stabilizing role for TGR5 would explain why bath application of its antagonist, SBI-115, reveals stochastic fluctuations in CIN firing that likely arise from both intrinsic ion channel activity and extrinsic synaptic inputs.

### 4.3. Bile Effects on CIN Inhibitory Current Rate and Amplitude

To further investigate the mechanisms underlying the excitatory effects of low BA concentrations (1 and 10 µM CA/DCA) on CIN firing, we examined the impact of 10 µM CA/DCA exposure on sEPSCs. Intracellular dialysis, which occurs during whole-cell patch-clamp recordings when the pipette solution mixes with the neuron’s cytoplasm, dilutes endogenous signaling molecules [29], which may on its own disrupt some signaling. Therefore, we selected a high bath concentration of BA (10 µM) during whole-cell recordings to ensure sufficient BA effects on CINs and account for possible changes in sensitivity. Although 10 µM CA/DCA exceeds the ∼2.5 µM peak brain concentration observed following ethanol or fatty food consumption, it reliably produced measurable effects while remaining within a physiologically informative range. Our results showed that 10 µM CA/DCA significantly suppressed sIPSCs, suggesting that low BA concentrations directly modulate GABAergic signaling. Interestingly, 10 µM CA/DCA also increased sIPSC amplitude despite reducing their frequency. This effect may be due to the partial inhibition of presynaptic GABA release, which would decrease the number of overlapping inhibitory events, reduce shunting from concurrent inputs, and thereby allow each remaining inhibitory event to generate larger postsynaptic currents.

Our experiments further demonstrated that 100 µM and 1 mM CA/DCA did not significantly affect sIPSC frequency, consistent with cell-attached recordings showing that these concentrations do not alter overall CIN firing. Notably, 1 mM CA/DCA effects on inhibitory currents was highly variable and maintaining the whole-cell seal was considerably more difficult compared to other BA conditions, causing several experiments to end early. These results are consistent with the known membrane-disrupting effects of BAs and may explain the inhibitory effects of 10 mM CA/DCA on CIN firing. Interestingly, although 100 µM and 1 mM CA/DCA did not change event frequency, both concentrations reduced the average sIPSC amplitude. The presence of amplitude alterations in the absence of frequency changes indicates that presynaptic release was likely unaffected, whereas postsynaptic mechanisms—such as receptor desensitization or subtle membrane alterations—may have contributed to these inhibitory effects.

Analysis of sIPSCs showed that, beyond exerting independent effects on event frequency and amplitude, bile also differentially modulates average current amplitude and the distribution of event amplitudes. At 10 µM, both the average amplitude and the occurrence of larger events increased, indicating selective enhancement of specific inputs via presynaptic facilitation and/or postsynaptic sensitization. At 100 µM, the average amplitude decreased without altering the amplitude size distribution, consistent with uniform postsynaptic dampening of all current amplitudes. At 1 mM, although the average amplitude decreased, the distribution shifted toward a higher occurrence of larger events, reflecting complex, non-uniform postsynaptic modulation—possibly arising from heterogeneous receptor clustering or differential effects on membrane microdomains that selectively enhance larger currents while reducing smaller ones. Together, these results highlight the nuanced role of BAs in inhibitory synaptic transmission.

### 4.4. Bile Effects on Inhibitory, SBI-115-Treated CIN Inputs and on Excitatory CIN Inputs

Cell-attached recordings revealed that the TGR5 antagonist SBI-115 abolished the excitatory effects that low BA concentrations (1-10 µM CA/DCA) have on CIN firing rates. However, when examining inhibitory currents, 10 µM CA/DCA continued to suppress inhibitory input even in the presence of SBI-115, indicating a TGR5-independent mechanism. This suggests that reduced inhibition contributes to, but does not fully account for, the excitatory effects of low BA concentrations— which likely involve additional signaling pathways that modulate CIN activity. To investigate whether one of these additional pathways involves modulation of excitatory transmission, we examined the impact of CA/DCA on excitatory CIN currents. Excitatory events were isolated by addition of 10 µM picrotoxin, a GABA_A_ and glycine receptor antagonist. CA/DCA did not significantly alter sEPSC frequency, indicating that BAs preferentially modulate GABAergic rather than glutamatergic inputs onto CINs, though these experiments are limited in scope since only one concentration was examined. Since sEPSCs appear largely unaffected by BAs at 10 μM, and SBI-115 does not block BA effects on sIPSCs, BAs may act directly on CINs, perhaps by modulating intrinsic excitability (e.g. via modulation of hyperpolarization-activated cation channels or Ca^2+^ activated K^+^ channels). In summary, while modulation of GABAergic transmission represents one mechanism by which BAs alter CIN activity, it does not fully explain the excitatory effects of low BA concentrations. Instead, BA effects on CIN activity likely result from the convergence of TGR5-dependent signaling on intrinsic channel activity, and additional TGR5-independent mechanisms. These TGR5-independent mechanisms may involve activation of other non–G-protein–coupled BA receptors, including FXRs, which are expressed in multiple brain regions [30], and S1PR2, which shows RNA expression in the NAc [31].

### 4.5. Direct and CIN-Mediated Bile Effects on DA Release in the NAc

Evidence of bile’s concentration-dependent effects on accumbal CIN firing prompted subsequent voltammetry experiments to examine potential indirect, CIN-mediated BA effects on accumbal DA release [2]. We found that 10 mM CA/DCA markedly decreased DA levels. Mediation analysis suggested that this inhibitory effect is primarily driven by suppression of CIN activity. However, DA release did not recover during the wash condition, indicating potential neuronal dysfunction from generalized membrane disruption at 10 mM CA/DCA. Such dysfunction could inflate the apparent indirect effect, making it appear that 10 mM CA/DCA inhibits DA release via CIN suppression, when it likely suppresses both CIN activity and DA release directly through membrane disruption.

Although low bile concentrations (≤10 µM CA/DCA) do not significantly alter average DA release over longer recordings, exposure to 1 and 10 µM CA/DCA in shorter, isolated experiments decreases variability and enables statistical detection of transient increases in DA release at 8 and 14 min, respectively. Mediation analysis indicates that the transient increase in DA release at 1 µM CA/DCA is primarily CIN-mediated. Despite the significant CIN-mediated promotion of DA release, the total bile effect on DA release remains largely nonsignificant. Interestingly, the direct effect of 1 µM CA/DCA on DA release, though nonsignificant, trends toward modest inhibition. Together, these findings suggest that subtle direct inhibition of DA release at low BA concentrations may counterbalance CIN-mediated stimulation, restricting DA increases to transient periods and preventing sustained changes in average DA levels.

In contrast to the effects observed at 1 µM CA/DCA, transient DA elevations during exposure to 10 µM CA/DCA showed no significant CIN-mediated contribution. The direct bile effect on DA activity at 10 µM CA/DCA was excitatory and nearly significant for the concentration response curve, while the total bile effect was excitatory and significant. These results suggest that transient DA increases at 10 µM CA/DCA are primarily driven by direct bile effects, with only minor contributions from CIN-mediated pathways. The nearly significant excitatory direct bile effect may involve the activation of BA receptors such as TGR5 and FXR [11]. Although the total bile effect is significant, it remains subtle—likely reflecting opposing inhibitory and excitatory mechanisms—which may explain the transient nature of DA release increases observed with 10 µM CA/DCA. Importantly, increased DA levels at 1-10 µM CA/DCA are not due to reduced DA clearance rates, as statistical analysis revealed no significant changes in reuptake or decay rates with BA exposure at these concentrations [21].

Taken together, our findings indicate that bile modulates DA release in the NAc through distinct, concentration-dependent mechanisms. At extreme supraphysiological concentrations (1–10 mM), bile suppresses DA release primarily via inhibition of CIN activity, although generalized membrane disruption likely contributes at this range. At physiological levels (1 µM), bile enhances DA release through CIN-mediated excitation, whereas at mildly supraphysiological concentrations (10 µM), bile promotes DA release predominantly through direct excitatory actions. These results suggest that BA concentrations induced in the brain by ethanol and fatty food consumption (∼1 µM) are sufficient to transiently increase DA release in the NAc, supporting the possibility that BAs may contribute to the reinforcing properties of these substances *in vivo*.

## 5. Conclusion

Fatty foods and ethanol stimulate hepatic, and possibly local brain, BA secretion, which may contribute to these substances’ reinforcing effects. The present study demonstrates that BAs modulate striatal activity *in vitro* and suggests that both hepatic- and brain-derived BAs may influence mesolimbic function *in vivo*. Our results show that BAs exert a concentration-dependent effect on accumbal CIN firing rates. We further demonstrate that low BA concentrations (1-10 µM CA/DCA) enhance CIN firing via at least two independent mechanisms: TGR5 activation and suppression of inhibitory CIN currents. Our whole-cell recordings suggest that BAs modulate CIN firing in a complex, concentration-dependent manner, selectively affecting event frequency, current amplitude, and the distribution of amplitude sizes. Additionally, our voltammetry recordings indicate that although low BA concentrations do not alter average DA release over time, exposure to 1 and 10 µM CA/DCA elicits transient increases in DA levels. These DA effects are likely mediated by both CIN-dependent mechanisms and direct BA actions on dopaminergic terminals in the NAc, and do not involve changes in DA clearance rates. Collectively, these findings offer compelling evidence for BA modulation of CIN activity and striatal DA signaling.

## 6. Statements and Declarations

Isabella Roque, Shimran Sharma, Philipp Mews, Summer L. Thompson, and Jordan T. Yorgason declare that they have no conflict of interest. All institutional and national guidelines for the care and use of laboratory animals were followed. This work was supported by the US National Institutes of Health (NIH) grant AA030577 to JTY. The authors have no relevant financial or non-financial interests to disclose. Material preparation, data collection, data analysis and figure creation were performed by Isabella Roque and Shimran Sharma. All authors contributed to the experimental design. The first draft of the manuscript was written by Isabella Roque and all authors commented on previous versions of the manuscript and figures. All authors read and approved the final manuscript and figures.

